# Pyroxasulfone resistance in *Lolium rigidum* conferred by enhanced metabolic capacity

**DOI:** 10.1101/116269

**Authors:** R Busi, A Porri, TA Gaines, SB Powles

## Abstract

**Author Contributions:** RB and AP designed and performed the experiments. RB, AP, TG, SP wrote the manuscript.

**One sentence summary:** This study provides novel insight into herbicide resistance conferred by GST-based detoxification to allow proactive intervention to minimize weed resistance evolution.

**Abstract:** The evolution of herbicide-resistant weed populations in response to synthetic herbicide selective pressure is threatening safe weed control practices achieved by these molecules. In Australia multiple-resistant populations of annual ryegrass (*Lolium rigidum*) are effectively controlled by soil-applied herbicides which provide adequate weed control.

In this study we define the mechanistic basis of the experimentally-evolved resistance to the soil-applied herbicide pyroxasulfone in a *L. rigidum* population. TLC and HPLC-MS provide biochemical confirmation that pyroxasulfone resistance is metabolism-based with identification and quantification of pyroxasulfone metabolites formed *via* a glutathione conjugation pathway in pyroxasulfone-resistant *L. rigidum* plants. The observed over-expression of two putative resistance-endowing *GST* genes is consistent with pyroxasulfone-resistance in parental plants (P6) and positively correlated to pyroxasulfone resistance in F_1_ pair-cross progenies. Thus, a major detoxification mechanism involves glutathione conjugation to pyroxasulfone and *GST* over-expression in pyroxasulfone-resistant *L. rigidum* plants. The definition of the genetic basis of metabolic resistance in weeds can be a first crucial step towards chemical means to reverse resistance and improve long-term weed resistance management.

## Introduction

In modern and mechanized agriculture, herbicide weed control is mandatory to avoid significant crop losses (Oerke, 2006). However, the evolution of adaptive traits conferring herbicide resistance in agricultural weeds is hampering the efficiency of weed control by herbicides (Beckie and Tardif, 2012; Powles and Yu, 2010).

Evolved herbicide resistance in weed species can be target-site-based, due to a nucleotide mutation changing a key amino acid substitution at a herbicide binding site (site of action) of a target enzyme structure Such target-Site Resistance (TSR) is usually single-gene inherited resistance (reviewed by Darmency, 1994; Dayan *et al.,* 2014; Délye, 2005; Kaundun, 2014; Tranel and Wright, 2002). Conversely, Non-Target-Site Resistance (NTSR) is often responsible for herbicide resistance. NTSR embraces any mechanisms that minimize herbicide injury by limiting toxic herbicide concentrations reaching herbicide sites of action. Important among NTSR mechanisms are constitutive enzymatic super families responsible for concerted secondary plant metabolism. Herbicide detoxification can schematically occur in four phases: phase I (oxidation), phase II (conjugation), phase III (transport) and phase IIII (further degradation/compartmentation) (reviewed by Délye *et al.,* 2013; Kreuz *et al.,* 1996; Yuan *et al.,* 2007). These enzymes can serendipitously mediate herbicide detoxification via herbicide metabolism and inactivation [e.g., cytochrome P450 mono-oxygenases (P450s), glutathione-S-transferases (GSTs; EC 2.5.1.18) or glucosyltransferases (GTs)] followed by herbicide sequestration (.e.g., ABC transporters) (Davies and Caseley, 1999; Edwards *et al.,* 2005; Hatzios and Burgos, 2004). Some herbicides that interact with a complex of primary targets (e.g., chloroacetamides which inhibit a complex system of elongases responsible for the biosynthesis of very long chain fatty acids, VLCFA) have only selected for NTSR mechanisms (reviewed by Busi, 2014).

The molecular definition of NTSR mechanisms is often complicated, for example P450s or GSTs are enzyme superfamilies with a multitude of gene family members that often interact as part of a shared ‘family business’ within a particular detoxification pathway (Yuan *et al.,* 2007). It has been shown that P450s can facilitate the oxidation or hydroxylation of a range of herbicide molecules (Werck-Reichhart *et al.,* 2000) and be responsible for herbicide metabolism in different crop species (e.g. maize, rice, wheat) and weeds (Gaines *et al.,* 2014; Iwakami *et al.,* 2014; Kreuz *et al.,* 1996). Glutathione-S-transferases (GSTs) are phase II enzymes that can allow herbicides metabolism through conjugation with the tripeptide glutathione (γ-glutamylcysteinylglycine) (Cummins *et al.,* 2009). Early studies on GSTs were conducted with crop plants to understand the basis of herbicide selectivity. For example, it was shown that expression levels of detoxifying GSTs in certain crops were much greater than in weeds to explain herbicide selectivity (Hatton *et al.,* 1996).

*Lolium rigidum* (Gaud.) is a genetically diverse, cross-pollinated weed species that is widespread in the southern Australian cropping system and has evolved resistance to many different herbicide modes of action (reviewed by Yu and Powles, 2014). In Australia the first selective herbicide deployed for *L. rigidum* control was the acetyl CoA carboxylase (ACCase)-inhibiting herbicide diclofop-methyl introduced in 1978, followed by the acetolactate synthase (ALS)-inhibiting herbicide chlorsulfuron in 1982. Heap & Knight (1986) reported the first case of cross-resistance to ACCase and ALS herbicides evolved by diclofop-methyl field selection. Currently, ACCase and ALS cross-resistance is widespread throughout the southern Australian cropping system (Malone *et al.,* 2013; Owen *et al.,* 2014).

In response to widespread ACCase and ALS herbicide resistance, there has been heavy reliance on pre-emergence soil-applied herbicides such as prosulfocarb, pyroxasulfone, triallate and trifluralin to which resistance currently remains at low levels (Busi and Powles, 2016; Powles *et al.,* 1988). The relatively new herbicide pyroxasulfone (VLCFA inhibitor) has become widely used in Australia, U.S.A and Canada. In Canada a recent study reported field-evolved resistance to pyroxasulfone and triallate in *A. fatua* (Mangin *et al.,* 2016). No field-evolved pyroxasulfone-resistant *L. rigidum* populations have thus far been identified, however we experimentally evolved pyroxasulfone resistance in *L. rigidum* by recurrent low-dose pyroxasulfone selection over a few generations (Busi *et al.,* 2012) and showed cross-resistance to the thiocarbamates prosulfocarb and triallate rapidly evolving in field collected populations (Busi and Powles, 2013, 2016). Here, we present studies to elucidate the mechanistic basis of pyroxasulfone resistance in *L. rigidum.*

## Material and Methods

### Plant material

#### Parental *L. rigidum* populations

The multiple resistant *L. rigidum* population SLR31 (hereinafter referred to as MR) evolved in the field following extensive herbicide selection. MR plants exhibit multiple herbicide resistance to different modes of action including the ACCase-inhibitor diclofop-methyl, the ALS-inhibitor chlorsulfuron (Christopher *et al.,* 1991), the mitosis inhibitor trifluralin (McAlister *et al.,* 1995), and the VLCFAE inhibitor S-metolachlor (Burnet *et al.,* 1994). This MR population is susceptible to pyroxasulfone (VLCFAE inhibitor) (Walsh et al 2011), prosulfocarb (VLCFAE inhibitor), and marginally resistant to triallate (Tardif and Powles, 1999). MR individuals were exposed to recurrent selection with below-label, sub-lethal doses of pyroxasulfone and experimentally evolved resistance to pyroxasulfone, prosulfocarb and triallate (Busi *et al.,* 2012; Busi and Powles, 2013). Progeny P6 was obtained by six consecutive cycles of recurrent herbicide selection consisting of pyroxasulfone selection at 60 g ha^-1^ (Progeny one, P1), followed by another pyroxasulfone selection at 120 g ha^-1^ (Progeny two, P2) 120 g ha^-1^ (Progeny three, P3) 240 g ha^-1^ (Progeny four, P4) then further subjected to two consecutive selections at 1000 (Progeny five, P5) and 2000 (Progeny six, P6) g prosulfocarb ha^-1^. The herbicide susceptible *L. rigidum* population VLR1 was the control in all experiments (hereinafter referred to as ‘S’).

### Herbicide assay

Herbicide survival response to pyroxasulfone in parental populations and F_1_ families grown in pots Viable seeds of *L. rigidum* populations P6, MR, S were germinated on 0.6% (v/w) solidified agar and planted into 2L pots containing commercial potting mixture (50% peatmoss, 25% sand and 25% pine bark) when the primordial root was visibly erupting from the seed coat. Approximately 2 hours after seeding the pots were treated with 0 (untreated), 25 or 100 g pyroxasulfone ha^-1^. For each herbicide dose there were four replicates (experiment 1), six replicates (experiment 2) or two replicates (experiment 3) with 25 viable germinated seeds treated per replicate. Survival was assessed in parental populations at 60 days after treatment (DAT) in experiment 1 prior to leaf material collection, 15 DAT in experiment 2 or 21 DAT in experiment 3 in F_1_ families in response to 100 g pyroxasulfone ha^-1^.

### Metabolism study

#### Chemical compounds

^14^C-labeled pyroxasulfone ([isoxazoline-3-^14^C]pyroxasulfone) synthesized by Amersham Biosciences Co., Ltd. (United Kingdom) with specific radioactivity of 1.7 MBq/m and > 99% puritywas used in this study. Pyroxasulfone (white powder, mp 130.7°C (degrees Celsius), water solubility at 20°C 3.49 mg/L, vp 2.4×10^-6^ Pa) and the synthetic compounds, 2-amino-5-[1-(carboxylmethylamino)-3-(5,5-dimethyl-4,5-dihydroisoxazol-3-ylthio)-1-oxopropan-2-ylamino]-5-oxopentanoic acid (M-15), 2-amino-3-(5,5-dimethyl-4,5-dihydroisoxazol-3-ylthio) propanoic acid (M-26) and 3-(5,5-dimethyl-4,5-dihydroisoxazol-3-ylthio)-2-hydroxypropanoic acid (M-29) were used. These compounds were synthesized by KI Chemical Research Institute Co., Ltd. (Japan) and their purities were > 98%.

#### Pyroxasulfone treatments

Pyroxasulfone treatments were performed with similar method reported by Tanetani *et al.* (2013). In brief, 13 *L. rigidum* pyroxasulfone-resistant P6 and -susceptible S plants were grown hydroponically up to the 4-leaf stage in 70 ml distilled water containing 70μl of liquid fertilizer containing 10% phosphoric acid, 6% nitrogen and 5% potassium (HYPONex, HYPONex JAPAN CORP., LTD.). The plants were then exposed to 1.3 ppm pyroxasulfone (approximately 3.3 μM). Four individual plants were harvested at three different time intervals corresponding to 1, 2 and 4 days after pyroxasulfone treatment and used for extraction and fractionation.

#### Extraction and fractionation

The methodology for extraction and fractionation of pyroxasulfone metabolites following pyroxasulfone treatment of *L. rigidum* plants is described in detail by Tanetani *et al.* (2013). In brief, following pyroxasulfone hydroponic treatment, *L. rigidum* plants were weighed, roots washed with 20 ml of acetonitrile and plants homogenized. Extraction of pyroxasulfone and its metabolites occurred in 150 ml of 25% acetone. The extracts were evaporated in vacuo and dissolved in 10 ml of 50% acetonitrile. The radioactivity of the extracts was measured with a liquid scintillation counter (LSC, TRI-CARB 2750TR/LL, PerkinElmer, United States). The radioactivity of the residues of the seedlings was measured with LSC after combustion by a sample oxidizer.

#### Metabolite identification

Pyroxasulfone and its metabolites were identified by comparison with standards, using thin layer chromatography (TLC) and LC-MS. For TLC analysis, an aliquot of each extract was applied to silica gel. The plates were firstly developed with a mixture of ethyl acetate/chloroform/methanol/formic acid (60/60/10/10, v/v/v/v) and secondly developed with a mixture of ethyl acetate/methanol/distilled water/formic acid (60/40/20/10, v/v/v/v). The subsequent determination of pyroxasulfone and its metabolites by TLC and LC-MS was performed as reported by Tanetani *et al.* (2013).

### RNA extraction and quantitative real-time PCR (qRT-PCR)

#### Experiment 1

Sixty days after pyroxasulfone treatment at 100 g ha^-1^ six resistant plants from the P6 population were identified and individually collected for total RNA extraction and q-PCR analysis. Similarly, six untreated individual plants (*n* = 6) from MR and S populations were individually harvested for the same q-PCR study, respectively. Two leaf segments of 2 cm were harvested from each individual 5-tiller plant and placed into a 25 mL tube. The individual plant represented the experimental unit as biological replicate.

#### Experiment 2

Fifteen days after pyroxasulfone treatment at 100 g ha^-1^ a total of 50 one-leaf surviving resistant P6 plants were harvested (2-cm plant tissue) and divided (*n* = 2) for total RNA extraction and subsequent q-PCR experiments. Also, 50 one-leaf plants emerging after pyroxasulfone treatment at 25 g ha^-1^ were harvested. In addition, 50 untreated MR, P6 and S one-leaf plants, respectively, were harvested for q-PCR analysis. Twenty leaf segments of 1 cm were harvested individually from 25 respective plants and pooled into a 25mL tube.

Total RNA was isolated from plant tissues by using RNAeasy extraction kit (Qiagen) and treated with DNA-free DNase (Ambion) to remove residual genomic DNA. One μg of total RNA was used for reverse transcription (Superscript III, Invitrogen) in a 20 μL volume reaction. Quantitative PCR was performed in a 384 well-plate using LightCycler 480 (Roche) and all reactions were conducted in three technical replicates and a negative control containing template and no primers for each amplification. 13 μL for each reaction included 6.5 μL of SyberGreen Master Mix, (SensiFAST), 0.25 μL of 0.5 pmol μL^-1^ primers, 3 μL of cDNA (diluted 1:10) and 3 μL of H_2_O. Reaction conditions were 3 min incubation at 95°C, 40 cycles of 95°C for 10 sec, 60°C for 20 sec, and 72°C for 10 sec followed by a melt-curve analysis to confirm single-product amplification.

Threshold-cycles (CTs) were obtained for each reaction using the Second Derivative Maximum method in the LightCycler 480 software (Roche). The mean of CT values for the three technical replicates for each sample was used to calculate the relative expression (RE) of the gene of interest using the following equation:

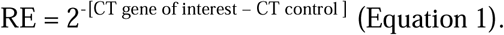

The control gene used in this assay was *isocitrate dehydrogenase* as described by Gaines et al. (2014). The relative expression of *GST-1* Tau class (contig 4546), *GST-2* Tau class (contig 5390), *GST-3* Phi class (contig 8676), *GST-4* Tau class (contig 13326), *GST-5* Phi class (contig 16302), and *P450-1 CYP72A* (contig 1604), *P450-2 CYP72A* (contig 2218), *P450-3 CYP716A* (contig 6783), *P450-4 CYP89A* (contig 6759) and *P450-5 CYP71B* (contig 12788) was quantified using primers described by Gaines et al., 2014 and shown in table S1.

### F_1_ pair-cross families for progeny test validation

Three parental resistant P6 plants (plant 1, 2 and 3) with the highest level of expression for contigs *GST-1* and *GST-2* (as determined in experiment 1) were vegetatively cloned (clone #1a, #2a and #3a) prior to being pair-crossed with three cloned plants (4, 5, and 6) from the parental population MR (clone #4a, #5a and #6a) and the other set of P6 (clone #1b, #2b and #3b) and MR cloned plants (clone #4b, #5b and #6b) were pair-crossed with two sets of three cloned plants (plant 7, 8 and 9) of the standard herbicide susceptible S (clone #7a, #8a, #9a and clone #7b, #8b, #9b, respectively). Thus 9 pair-crosses were established as [F1 #1 (1a × 4a)], [F1 #2 (1b × 7a)], [F1 #3 (4b × 7b)], [F1 #4 (2a × 5a)], [F1 #5 (2b × 8a)], [F1 #6 (5b × 8b)], F1 #7 (3a × 6a)], [F1#8 (3b × 9a)] and [F1 #9 (6b × 9b)] and the seed progeny was individually collected from each mother plant and identified as 18 distinct F_1_ families.

#### Experiment 3

As described above, the seed from all F_1_ families obtained by pair-crosses was treated with 100 g pyroxasulfone ha^-1^ to determine the correlation between sum of expression levels of *GST-1* and *GST-2* measured in parental plants resistant P6, MR and S and the herbicide response of those generated F_1_ seed progenies as the result of a pair-cross.

### Statistical analysis

For all the *L. rigidum* populations analysed in this study graphical data relative to the resistance phenotype are presented as percent (%) of seed germination and seedling survival or gene expression relative to population S set as equal to 1. Two main types of analysis were conducted to compare and separate population mean values for survival and gene expression levels. Comparisons among survival rates were assessed by chi-square (χ^2^) heterogeneity test performed using the statistical software *R* (version 3.02) with the command *prop.test.* Relative gene expression were subjected to ANOVA and population means (P6 vs. MR vs. S) separated by Tukey’s HSD (α = 0.05). Pearson’s correlation coefficient (r), 95% confidence intervals and two-tailed *P* values for pair wise combination of *GST-1* and *GST-2* expression levels in parental plants and plant survival (%) in the F_1_ seed progeny in pair-crosses (P6 × MR × S) was calculated with GraphPad Prism (GraphPad Software, Inc. La Jolla, CA 92037 USA).

## Results

### Response to pyroxasulfone treatments of resistant P6, MR and S L. rigidum plants prior to molecular analysis

When treated at the recommended dose of pyroxasulfone (100 g ha^-1^) there was 54% survival of the resistant P6 plants. As expected, for the parental MR and the standard herbicide-susceptible S populations there was only 5% survival (Figure 1A). Surviving resistant P6 plants 60 DAT were then used for the subsequent molecular analysis and compared to untreated MR and S plants. The herbicide assays were repeated with 48% plant survival observed in P6 plants (data not shown). A similar level of herbicide stress in MR and S plants was obtained at the lower dose of 25 g pyroxasulfone ha^-1^ (45% and 27% survival, respectively, data not shown). From this dose-response study (experiment 2) 50 1-leaf emerging seedlings 15 DAT were bulk collected for each resistant P6, MR and S *L. rigidum* population and subjected to molecular analysis.

**Fig 1.**
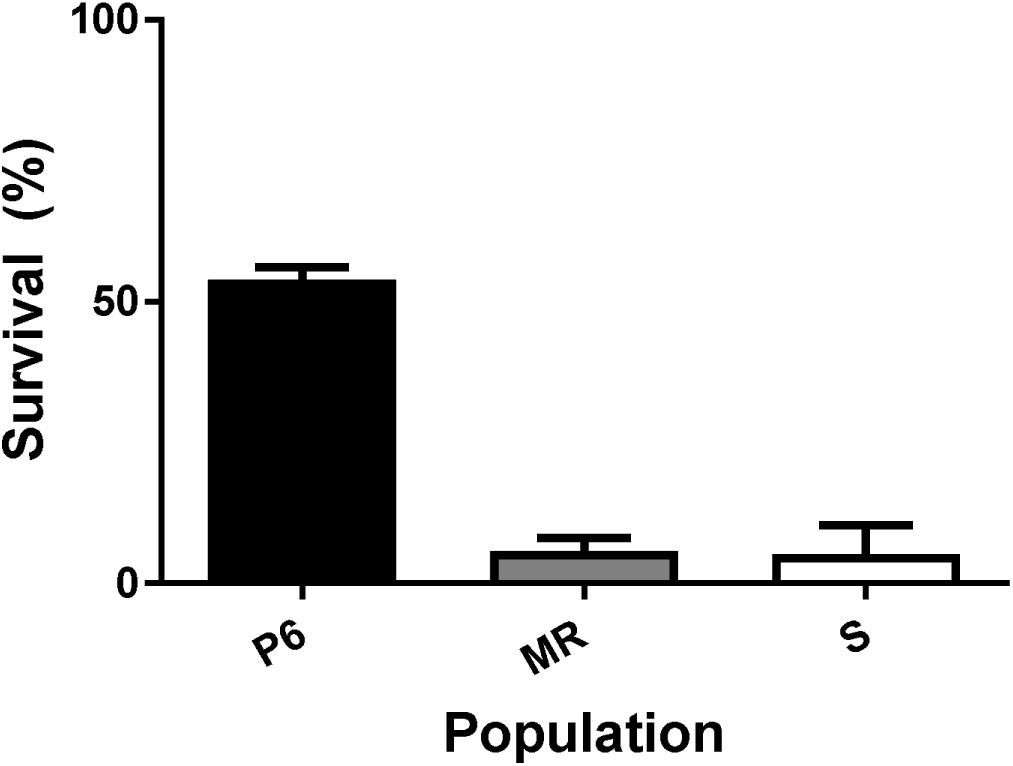
Mean plant survival (%) as ratio of actively growing seedlings versus seeds treated ± standard errors (SE) in pyroxasulfone treated *Lolium rigidum* plants. Survival ± SE (*n* = 4) assessed as seedling emergence in pot cultured plants assessed 60 days after 100 g pyroxasulfone ha^−1^ treatment in pyroxasulfone-resistant progeny P6 (black bar), parental MR (grey bar), herbicide susceptible S population (white bar) or wheat (W, light grey bar).

### ^14^C-pyroxasulfone metabolites analysis in pyroxasulfone-resistant P6 L. rigidum plants

Following root application of ^14^C-pyroxasulfone to *L. rigidum* plants at the 3-leaf stage, the total radioactivity was determined over time. Pyroxasulfone-resistant P6 plants absorbed from 8% (1 DAT) to 25% (4 DAT) of pyroxasulfone provided, corresponding to a concentration of 10.71 μg eq./g plant tissue harvested (Table 1). Similar results were obtained in pyroxasulfone-susceptible (S) *L. rigidum* and wheat plants as described by Tanetani *et al.* (2013).

**Table 1.** Amount of radioactivity in plant *L. rigidum* plants treated with [isoxazoline-^14^C] pyroxasulfone and harvested 1, 2 and 4 days after treatment.

| Population |  | Days after treatment | Plant fresh mass (g) | Radioactivity (µg eq.)* |  |  | Recovery (%) | Concentration(µg (µg eq./g)** |
| --- | --- | --- | --- | --- | --- | --- | --- | --- |
|  |  |  |  | Extraction | Residue (not extracted) | Total radioactivity |  |  |
| P6 |  | 1 | 1.51 | 5.88 | 0.88 | 6.76 | 8 | 4.48 |
| P6 |  | 2 | 1.76 | 10.00 | 1.12 | 11.13 | 12 | 6.33 |
| P6 |  | 4 | 2.08 | 20.00 | 2.06 | 22.06 | 25 | 10.71 |
| S<sup>+</sup> |  | 1 | 1.82 | 3.5 | 0.6 | 4.1 | 4 | 2.3 |
| S<sup>+</sup> |  | 2 | 1.89 | 5.3 | 0.6 | 5.9 | 6 | 3.1 |
| S<sup>+</sup> |  | 4 | 1.87 | 8.2 | 0.6 | 8.8 | 10 | 4.7 |

\*Values are expressed as the amount equivalent to pyroxasulfone.
\*\* Concentration is the amount of parent compound equivalent (µg eq.) to plant fresh weight (g).
<sup>++</sup> Data of S plants are shown in Table 2 (Tanetani *et al.*, 2013).

The total radioactivity absorbed in the resistant P6 plants was approximately 2-fold higher than in S plants (Table 1). Equally, the total amounts of the metabolites found up to four days after treatment (DAT) in the resistant P6 plants were larger than in S plants. The parental ^14^C-pyroxasulfone was rapidly degraded into several metabolites. The decomposition rate of ^14^C-pyroxasulfone in the P6 pyroxasulfone resistant plants was much greater and up to 4-fold higher at 1 DAT than in the S plants. In resistant P6 plants at 1 and 2 days after pyroxasulfone treatment, the ratio of pyroxasulfone in R biotype was lower than that of S biotype, and the ratio of metabolites in the R biotype was higher than in the S biotype (Table 2).

**Table 2.** Proportion of absorbed parental pyroxasulfone (%) and its metabolites (identified and unknown) in *L. rigidum* plants treated with pyroxasulfone and harvested 1, 2 and 4 days after treatment.

| Population | Days after treatment | Pyroxasulfone (µg eq./g) | Metabolites identified |  |  | Total (identified) | Metabolites unknown |  |  |  |  |  | Total |
| --- | --- | --- | --- | --- | --- | --- | --- | --- | --- | --- | --- | --- | --- |
|  |  |  | M-26 | M-29 | M-29-glc |  | Uk-1 | Uk-2 | Uk-3 | Uk-4 | Others |  |  |
| P6 | 1 | 12.2 (0.57)* | 21.8 | 7.8 | 15.7 | 45.3 (1.76) | 5.2 | 6.1 | 9.1 | 5.2 | 0.9 |  | 84.0 |
| P6 | 2 | 4.5 (0.28)* | 22.5 | 17.1 | 11.7 | 51.3 (2.92) | 3.7 | 5.4 | 12.6 | 3.6 | 6.8 |  | 87.9 |
| P6 | 4 | 4.6 (0.49)* | 20.0 | 12.7 | 20.0 | 52.7 (5.07) | 2.7 | 4.6 | 11.8 | 2.7 | 6.6 |  | 85.7 |
| S<sup>+</sup> | 1 | 46.4 (1.07) | 13.7 | 3.6 | 4.6 | 21.9 (0.50) | 4.6 | 1.8 | 2.7 | 3.6 | 4.4 |  | 85.4 |
| S<sup>+</sup> | 2 | 26.4 (0.82) | 18.2 | 8.2 | 10.0 | 36.4 (1.13) | 8.2 | 1.8 | 3.6 | 6.4 | 7.0 |  | 89.8 |
| S<sup>+</sup> | 4 | 9.1 (0.43) | 24.6 | 10.0 | 13.7 | 48.3 (2.27) | 8.2 | 1.8 | 8.2 | 7.3 | 10.3 |  | 93.2 |

\*\* Values in parentheses indicate concentration (µg eq./g). <sup>++</sup> Data of S plants are shown in Table 3 (Tanetani *et al.*, 2013).

In the extracts from resistant P6 and S plants, a total of eight metabolites were evident in TLC analysis (Figure 2). Six of these metabolites (TLC spots), namely pyroxasulfone, Uk-1, Uk-3, M-26, M-29 and glucose conjugate of M-29 (M-29-glc) were the same chemical compounds as those detected in wheat (Figure 2, Table 2) (Tanetani *et al.,* 2013). Considering the ratio of the radioactivity of each metabolite, M-26, M-29, and M-29-glc were the main metabolites identified in wheat and the S *L. rigidum.* M-26 was generated by liberating glutamic acid and glycine from glutathione conjugate of the isoxazoline ring (M-15) and M-26 was metabolized to M-29 by oxidative deamination. Subsequently, M-29-glc was generated by glucose conjugation of M-29. These metabolic processes indicated that the main metabolites (M-26, M-29 and M-29-glc) are assumed to be formed *via* glutathione conjugation of the isoxazoline ring of pyroxasulfone. Thus the main route of pyroxasulfone metabolism appears to be the cleavage of methylensulfonyl linkage by glutathione-conjugation of the isoxazoline ring (Tanetani *et al,* 2013).

**Fig 2.**
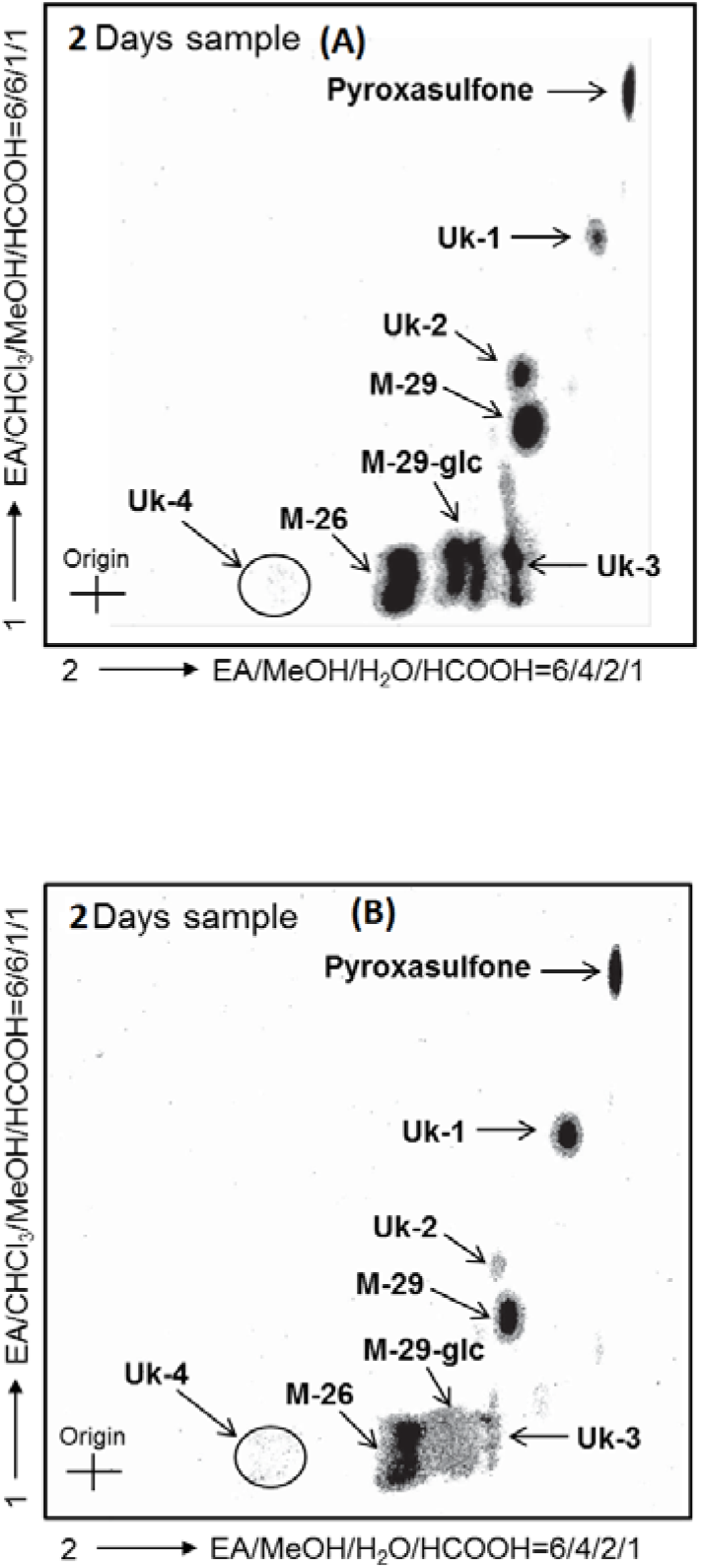
Two dimensional TLC of the extract from R biotype of rigid ryegrass after treatment with ^14^C-pyroxasulfone (4 DAT) in (A) pyroxasulfone-resistant (P6) versus (B) pyroxasulfone-susceptible (S) *L. rigidum* plants. ^++^ Figure 2B reports data shown in Figure 1 of (Tanetani et al., 2013).

### Transcript levels of genes encoding herbicide-metabolizing enzyme in resistant P6, MR and S L. rigidum plants

To assess whether pyroxasulfone resistance is associated with increased transcript levels of herbicide-metabolizing genes, the expression levels of five putative *P450s* and *GSTs* previously identified in resistant *Lolium* populations (Gaines et al., 2014) were determined by quantitative real time PCR. The tested *P450s* and *GSTs* were named from 1 to 5 (see material and methods). In this assay the P6 pyroxasulfone resistant individuals were compared with the untreated susceptible MR individuals and susceptible S individuals (VRL1). The transcript quantification was performed on six different biological replicates and the statistical significance among the different individuals was assessed using Tukey’s HSD and ANOVA tests. The mRNA level of *P450-1* was increased around 6 and 4 times in both R P6 individuals and S MR individuals compared with the S plants, respectively (P < 0.01) (Figure 3). However, there was no significant difference in *P450-1* expression in resistant P6 compared to MR individuals. The mRNA abundances of *P450-2, P450-4* and *P450-5* were not significantly different among resistant P6, MR and S plants, while the expression of *P450-3* was 5- and 3-fold reduced in resistant P6 and MR, respectively, compared with S plants (P < 0.01) (Figure 3). The transcript levels of *GST-1* were around 9 times higher in R P6 individuals compared to both MR and S plants. Likewise, the mRNA levels of *GST-2* were around 6 and 3 times more abundant in R P6 plants compared to MR and S individuals, respectively (Figure 4). The upregulation of these two *GSTs* was consistently found in all tested P6 biological replicates. Tukey’s multiple comparisons test of *GST-1* and *GST-2* expression data showed high statistical significance (*p* value ≤ 0.01). In contrast, the expression levels of *GST-3, GST-4* and *GST-5* were not significantly different among resistant P6, MR and S individuals (Figure 4). Thus, in the resistant P6 plants the increased transcript levels of *GST-1* and *GST-2* are associated with pyroxasulfone resistance. For further confirmation the expression levels of these two *GTSs* were quantified in resistant P6, MR and S one-leaf stage plants, 15 days after pyroxasulfone pre-emergence treatment. Resistant P6 individuals were treated with 100 g pyroxasulfone ha^-1^ whereas MR and S plants were treated with a sub-lethal 25 g ha^-1^. In addition, to assess whether the expression of *GST-1* and *GST-2* is constitutively increased in the resistant P6 plants independently of the herbicide treatment, untreated resistant P6, MR and S individuals were also collected. The transcript levels of *GST-1* and *GST-2* in untreated resistant P6 plants were significantly (*p* value ≤ 0.01) higher than in MR and S plants, with a calculated 7- and 4-fold higher relative gene expression, respectively. Similar results indicating *GST1-1* and *GST-2* over-expression were found in the pyroxasulfone treated plants (Figure 5).

**Fig 3.**
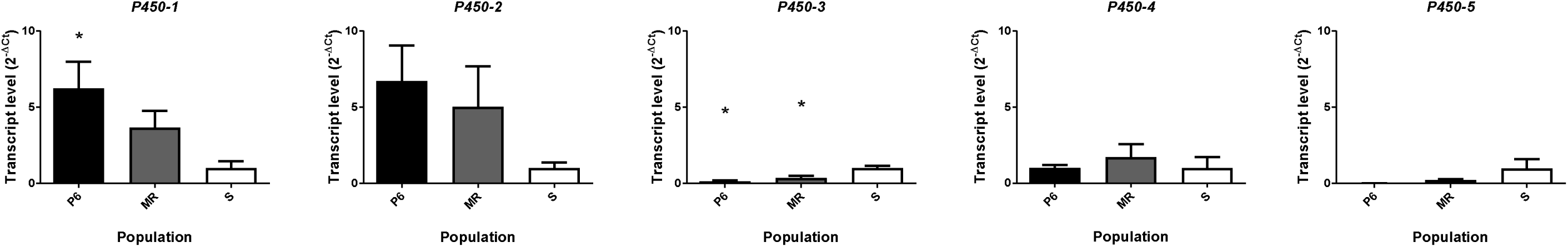
Transcript levels of *P450* genes in *L. rigidum* plants harvested at the 5-tillers stage sixty days after 100 g pyroxasulfone ha^-1^ treatment in pyroxasulfone-resistant progeny P6 (black bars), untreated parental MR population (grey bars) or herbicide untreated susceptible S population (white bars). Transcript levels were assessed by real-time RT-PCR and *Isocitrate dehydrogenase* was used as internal control gene. Transcript abundance (gene expression) was normalized to the level of the S population. Data shown are means of six biological replicates (±standard error) [* indicate significant difference to the S population (treated or untreated) after ANOVA analysis and *post-hoc* Tuckey test P < 0.01].

**Fig 4.**
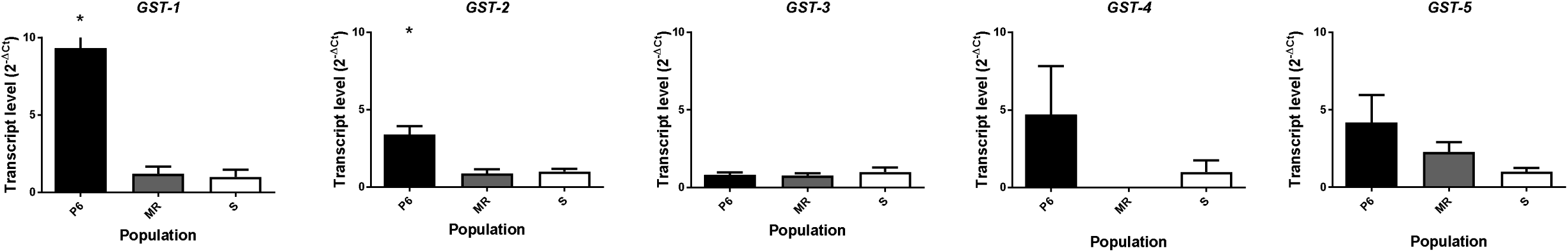
Transcript levels of *GST* genes in *L. rigidum* plants harvested at the 5-tillers stage sixty days after 100 g pyroxasulfone ha^−1^ treatment in pyroxasulfone-resistant progeny P6 (black bars), untreated parental MR population (grey bars) or herbicide untreated susceptible S population (white bars). Transcript levels were assessed by real-time RT-PCR and *Isocitrate dehydrogenase* was used as internal control gene. Transcript abundance (gene expression) was normalized to the level of the S population. Data shown are means of six biological replicates (±standard error) [* indicate significant difference to the S population after ANOVA analysis and *post-hoc* Tuckey test P < 0.01].

**Fig 5.**
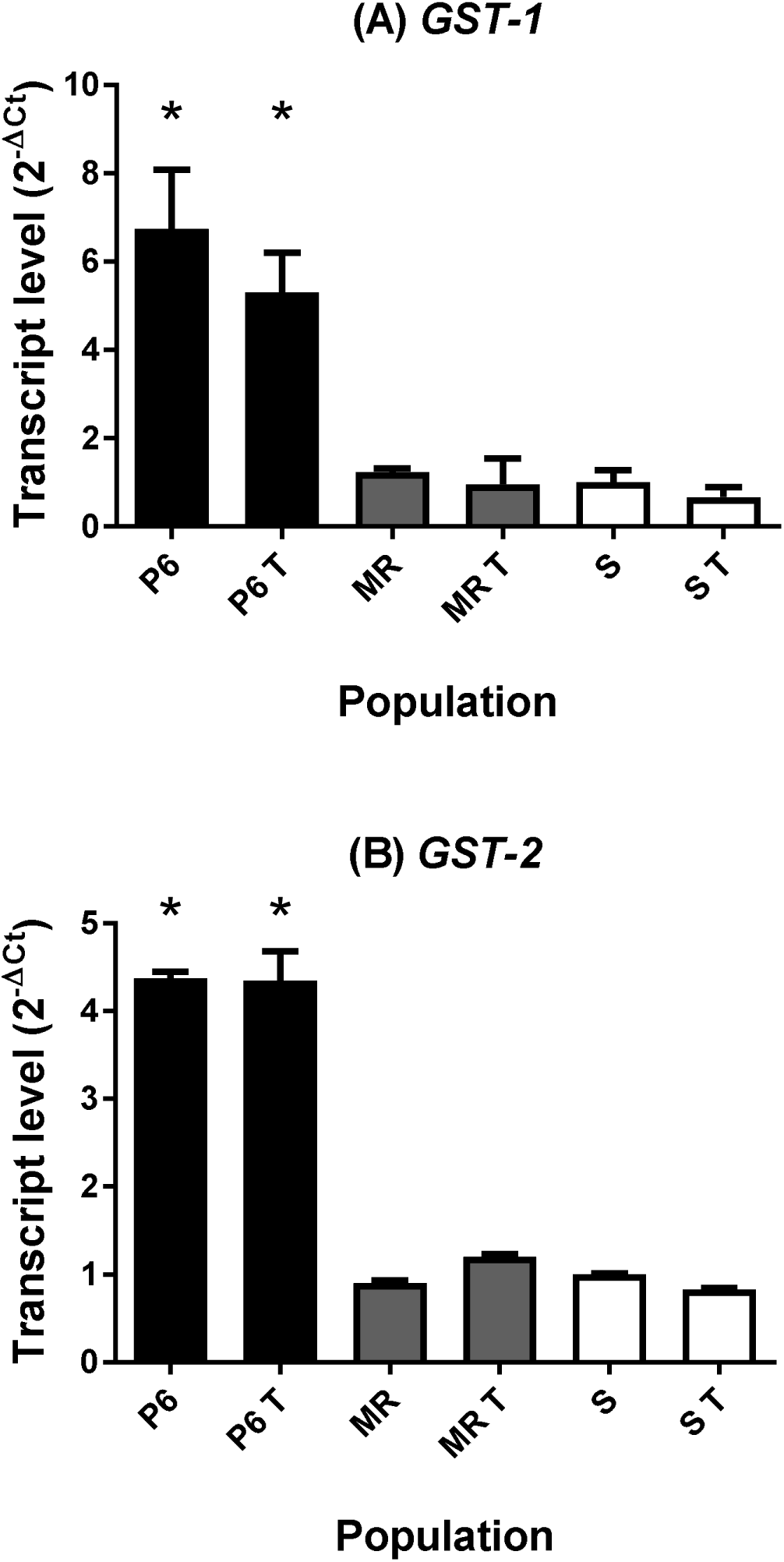
Transcript levels of *GST-1* (A) and *GST-2* (B) genes in one leaf stage *L. rigidum* plants harvested fifteen days after the application of 100 g pyroxasulfone ha^−1^ treatments (T) versus untreated P6 plants (black bars), treated (25 g pyroxasulfone ha^−1^) or untreated parental MR individuals (grey bars) or treated (25 g pyroxasulfone ha^−1^) or untreated susceptible S plants (white bars).

### Correlation between GST expression and pyroxasulfone resistance in F_1_ families

We assessed phenotypic pyroxasulfone resistance in 18 individual F_1_ families (reciprocal pair crosses) generated with three cloned resistant P6, MR and S plants. The mean plant survival assessed in F_1_ families generated *via* pair-cross of resistant P6 with MR was significantly higher than that of F_1_ families obtained with pair-cross of P6 with S (χ^2^ = 17; P < 0.001) which respectively was greater than survival in F_1_ from MR with S crosses (χ^2^ = 16; P < 0.001) (Figure 6). A positive and significant correlation (P < 0.001) was found between the sum of expression levels of *GST-1* and *GST-2* in parental plants and survival in F_1_ families with a calculated Pearson coefficient of *r* = 0.698 (data not shown) (Figure 6).

**Fig 6.**
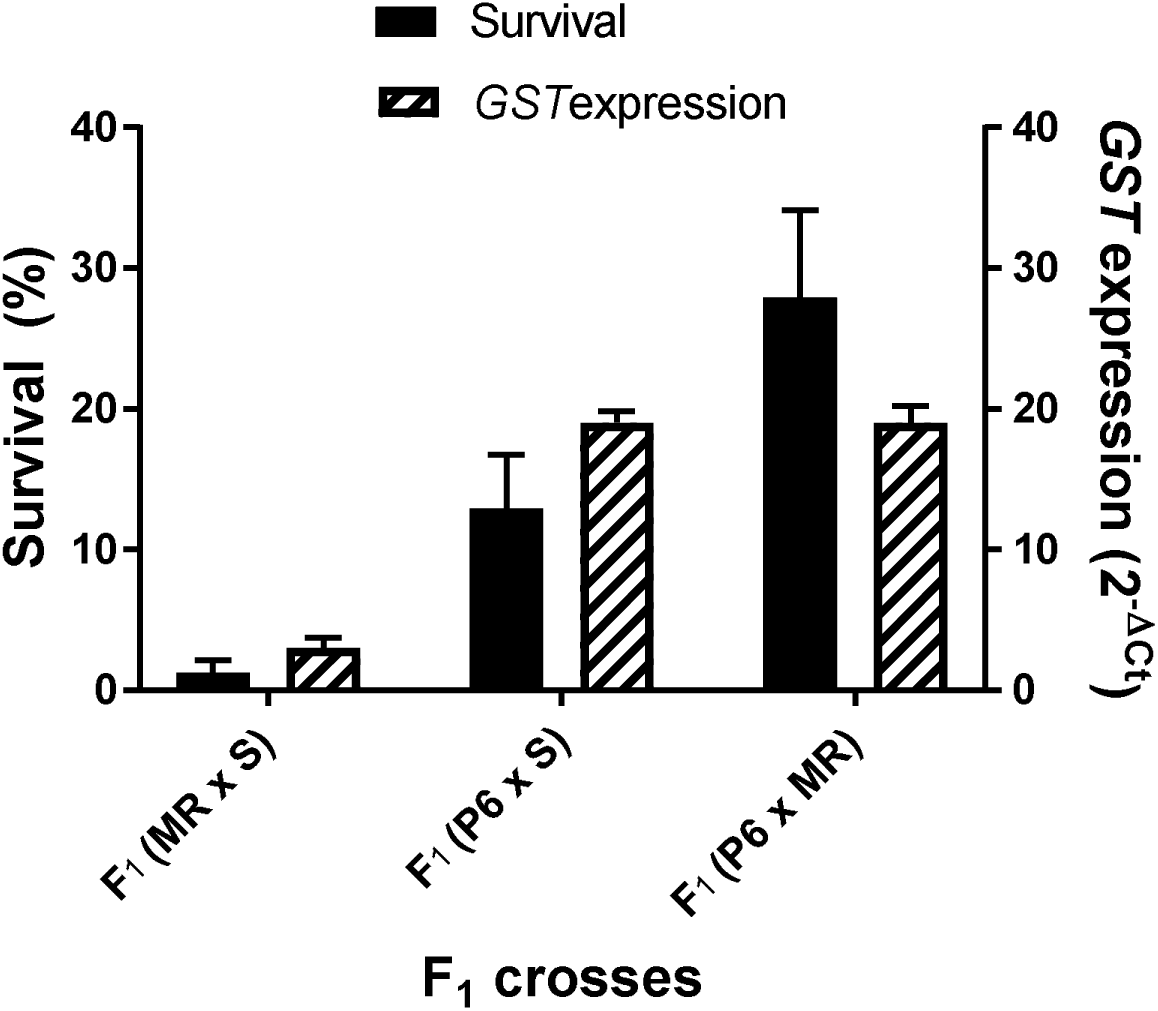
Mean plant survival (black bars, *n* = 6 F_1_ families obtained from 6 individual parental plants used in 3 pair crosses) of three types of F_1_ families (MR × S; P6 × S and P6 × MR) and sum of *GST-1* and *GST-2* expression (striped bars, *n* = 6 parental plants used in 3 pair crosses) to understand the correlation between the sum of *GST* expression levels of parental plants and plant survival observed in a total of 18 F1 families. Bars are mean values ± SE.

## Discussion

### Contribution of GST-1 and GST-2 to pyroxasulfone resistance in the P6 population

This study represents a major step towards characterizing the definition of the biochemical and genetic basis of pyroxasulfone resistance. Both metabolic and gene expression data supports a causative role for GST-mediated pyroxasulfone-glutathione conjugation. The mechanistic basis for pyroxasulfone resistance in the P6 resistant *L. rigidum* is metabolism-based with resistant plants displaying enhanced capacity to detoxify pyroxasulfone *via* a glutathione conjugation pathway. This result was clearly evident in resistant P6 plants one day after pyroxasulfone treatment, with approximately 85% of pyroxasulfone metabolized into several different metabolites. The combined TLC and LC-MS work indicates that the metabolites formed in pyroxasulfone-resistant P6 plants were likely formed *via* GSTs catalysing glutathione conjugation of the isoxazoline ring of pyroxasulfone and then the subsequent production of three main metabolites. A previous study reported the same metabolic pathway to explain the much greater metabolic detoxification of ^14^C-pyroxasulfone in tolerant wheat plants relative to pyroxasulfone-susceptible *L. rigidum* to explain safety versus toxicity in crops versus grass weeds (Tanetani *et al.,* 2013).

Since first reported to endow resistance to thiocarbamate herbicides (Lay and Casida, 1976) it has become clear that plant GSTs can catalyze conjugation of the tripeptide glutathione (γ-glutamyl-cysteinyl-glycine; GSH) with certain herbicides (Dixon *et al.,* 2002). The GST enzyme superfamily includes two plant specific classes [Phi (F) and Tau (U)] associated with herbicide resistance in weeds (Cummins *et al.,* 2011). Thus, the electrophilic nature of some herbicide molecules, often after initial P450-mediated hydroxylation, can bind to the cysteine residue of glutathione as the first step in this detoxification pathway (Fuerst, 1987). These chemical reactions involving K_3_ herbicides and GSH are similar to the covalent binding of the KCS enzymatic complex identified as one of the primary target for these VLCFAE-inhibiting herbicides (Böger *et al.,* 2000; Eckermann *et al.,* 2003). Crop selectivity to several different chloroacetamide herbicides is similarly mediated by enhanced GST activity (Lamoureux and Rusness, 1989; Leavitt and Penner, 1979). Thus, as pyroxasulfone levels decreased at a much faster rate in pyroxasulfone-resistant *L. rigidum* and tolerant wheat plants than in susceptible *L. rigidum* plants, this study suggest similarities in metabolic detoxification of pyroxasulfone between wheat and pyroxasulfone-resistant *L. rigidum.*

This study provides evidence that a significant increase in constitutive *GST* gene expression is associated with pyroxasulfone resistance at the individual parent plants and at the population level, as both *GST-1* and *GST-2,* both Tau class, had significantly higher transcription in P6 individuals than in MR or S individuals. We did not observe any additional upregulation of *GST-1* and *GST-2* in pyroxasulfone-treated versus untreated individuals indicating that in our P6 individuals the over-expression of these herbicide-metabolizing genes is constitutive. In a previous inheritance study we showed that pyroxasulfone resistance in *L. rigidum* was likely governed by semi-dominant allele(s) segregating at one major locus (Busi *et al.,* 2014). Importantly, here we provide evidence with a progeny test that GST overexpression in parental plants correlates with plant survival in F1 progenies. Thus, major traits for pyroxasulfone resistance evolved by recurrent herbicide selection of *L. rigidum* individuals (Busi *et al.,* 2014; Busi *et al.,* 2012) are now likely associated with greater constitutive expression of certain GST genes. The upregulation of two different genes in a trait inherited as a single semi-dominant allele could be explained if *GST-1* and *GST-2* were closely linked on the same chromosome, thereby producing an inheritance pattern consistent with a single locus. Another possibility is that transcription of the two different genes may be co-regulated by a single transcription factor, which would also produce a single gene inheritance pattern. In wheat plants *GST* (*TaGSTU4*) over-expression induced by the safener fenchlorazole-ethyl was found to mediate resistance to the ACCase-inhibiting herbicide fenoxaprop-ethyl and the K3 herbicide dimethenamide (Thom *et al.,* 2002). BLAST analysis reveals high similarities between *TaGSTU4* and *GST-1* (contig score 205, E-value 1.6 9 10^-53^) (Gaines *et al.,* 2014). Similarly, there was increased expression of *GST-5,* (Phi class, contig 16302 with 94.5% similarity in 145 bp) as reported by (Gaines *et al.,* 2014) to the *LrGSTF1* homologue of *AmGSTF1* endowing fenoxaprop-ethyl resistance in *A. myosuroides* (Cummins *et al.,* 2013) in both parental MR and pyroxasulfone-resistant P6 plants. Thus, this specific *GST-5* could confer some level of pyroxasulfone-resistance, although the available experimental evidence is not fully compelling. Other studies on transcriptome analysis provide additional evidence of *GST* over-expression conferring metabolic herbicide resistance in French populations of the grass weed *Lolium* (Duhoux *et al.,* 2017; Duhoux *et al.,* 2015; Gaines *et al.,* 2014). In *L. rigidum,* specific resistance to specific herbicides may be conferred by specific cytochrome P450s (Yu and Powles, 2014). For example, enhanced herbicide metabolism was shown in *L. rigidum* and wheat plants in response to the same ALS-inhibiting herbicide with evidence that resistance was likely mediated by cytochrome P450 (Christopher *et al.,* 1991). In a recent study we provided evidence of partial pyroxasulfone resistance reversal (approx. 40%) with the use of the organophosphate insecticide phorate which is believed to inhibit herbicide detoxifying mechanisms such as cytochrome P450 enzymes (Busi *et al.,* 2016). However, this study shows that the expression levels of the five tested *P450s* did not substantially differ among P6 and MR individuals as *P450-1* (*CYP72A*) and *P450-2* (*CYP72A*) were both up-regulated in pyroxasulfone-resistant P6 plants as well as parental pyroxasulfone-susceptible MR plants. Thus, evidence of enhanced rates of P450 activity conferring resistance to specific K3 herbicides such as pyroxasulfone in *L. rigidum* remains not fully compelling or understood.

Taken together, these data with the above experiments suggest that the increased transcription of *GST* constitutively occurs at crucial developmental stages in pyroxasulfone-resistant *L. rigidum* individuals. Further foundation work remains to be done starting from *de novo* transcriptome assembly and comparative transcriptomics analysis to unravel patterns of selection, mechanisms, gene expression and gene interactions driving the evolution of multiple resistance in major grass weeds.

## Acknowledgements

The AHRI Team is a center of excellence for the study of herbicide resistance funded by the Grains Research Development Corporation. The partial funding from Kumiai Chemical Industry, the Australian Research Council (LP0882758) and Bayer CropScience (Grants4Target #2016-2-22) is acknowledged.

